# Effect of real-time video feedback on the gait foot placement regularity and symmetry in individuals with trans-tibial amputation: A case study

**DOI:** 10.1101/381996

## Abstract

Low stability and asymmetrical gait are two important challenges in gait of individuals with trans-tibial amputation. The aim of this case report was to determine the feasibility and effectiveness of a real-time video feedback of foot placement on stability (i.e. foot placement regularity) and symmetry of trans-tibial amputees. Furthermore, we aimed at differentiating the effect of video feedback from front, back and lateral views on amputees’ walking. Seven male participants with unilateral trans-tibial amputation took part in this study. Each participant walked for 2 minutes in four conditions: no video feedback, video feedback from front view, video feedback from back view and video feedback from lateral view. Participants’ motion was recorded using five motion capture cameras. Nonlinear measure of sample entropy was adopted to quantify foot placement regularity (i.e. stability). The symmetry of step length and width was also determined using symmetry angle algorithm. Results demonstrated that video feedback significantly led to more regular and more symmetrical step length. Our findings suggest that amputees rely on visual feedback to have more stable and more symmetrical gait. The findings of this study suggest that implementing of video feedback gait training could reduce neuromusculoskeletal problems resulted from asymmetrical gait.

## 1. INTRODUCTION

It has been reported that people with lower limb amputation exhibit stability problems and are at higher risk of falling compared to nondisabled people (Hak et al., 2013; Lamoth, Ainsworth, Polomski, & Houdijk, 2010). This might be due to disruption of amputees’ motor and proprioceptive functions implying that people with amputation might rely more on visual feedback to maintain stability both in standing (Ku, Abu Osman, & Wan Abas, 2014) and walking (Beurskens, Wilken, & Dingwell, 2014). In addition, recent studies have calculated the step and/or stride regularity as a method of evaluating gait stability (Lamoth et al., 2010; Parker, Hanada, & Adderson, 2013). Considering together, it could be implied that by modifying foot placement regularity, it might be possible to increase walking stability.

In addition to stability, amputees have also asymmetrical gait (e.g., asymmetrical step length or step width). It is mainly because they are not able to (or prefer not to) put weight on the remaining parts of the amputated limb (i.e. residual limb) which results in an asymmetrical gait (Nolan et al., 2003). Gait asymmetry may lead to various neuromusculoskeletal problems such as pain and osteoarthritis in the intact limb as well as osteopenia/osteoporosis in the residual limb (Ephraim, Wegener, Mackenzie, Dillingham, & Pezzin, 2005; Kaufman, Frittoli, & Frigo, 2012). The correction of foot placement asymmetry in amputees could thus be beneficial to reduce the risk of orthopedic complications and thus to facilitate performing of everyday tasks.

It has been reported in several studies that gait training using visual feedback is a successful training protocol for people with different disorders such as Parkinson’s disease, stroke, trans-tibial amputation and total hip arthroplasty (Hamacher, Bertram, Folsch, & Schega, 2012). For instance, it has been shown in a study that a real-time visual feedback decreased energy consumption of amputees during treadmill walking (Davis et al., 2004). Darter and Wilken (2011) have shown that using virtual reality (VR) gait training improved frontal plane hip, pelvis, and trunk motion. In another study, Cho and Lee (2014) demonstrated that visual feedback gait training affected dynamic balance and gait in chronic stroke patients. Using VR with perturbation training, studies also demonstrated that amputees’ gait improved with VR and perturbation training (Kaufman, Wyatt, Sessoms, & Grabiner, 2014; Sheehan, Rabago, Rylander, Dingwell, & Wilken, 2016). However, there is no study focusing on realtime visual feedback of foot placement (i.e., step length and width) regularity and asymmetry in people with amputation. The implication being that if visual feedback increases foot placement regularity (and thus stability) and symmetry in amputee’s gait, then it can be a feasible and effective training method to improve gait stability and symmetry in the amputees. Another issue which has been neglected in previous studies is the direction of visual feedback. The studies using VR systems only provide a back view of the participants, which could be due to its intuitive nature as well as the limitation of the VR systems. Nevertheless, it is possible to provide visual feedback to participants from front, back and lateral views and currently it is not clear which direction affects participants’ stepping regularity and symmetry. Finally, the VR systems are expensive systems the application of which in small or medium size clinical settings (e.g., physical therapy clinics) might be impossible. An alternative could be video feedback systems which are easy to implement and also affordable.

The primary aim of this case report was therefore to determine the feasibility and effectiveness of a real-time video feedback of foot placement on foot placement regularity and symmetry of trans-tibial amputees. Moreover, this study aimed at differentiating the effect of video feedback from front, back and lateral views on amputees’ walking. We hypothesized that real-time video feedback of foot placement could regulate foot placement regularity and symmetry. In addition, it was hypothesized that all three front, back and lateral view video feedbacks would affect participants’ walking.

## 2. METHODS

### 2.1. Participants

Seven male participants with unilateral trans-tibial amputation took part in this study (Table 1). Inclusion criteria for participation in the study were: 1) being active community ambulators based on their self-report, 2) having normal visual ability, 3) having no orthopedic or neurological impairments other than amputation that might affect their walking ability (examined by a physical therapist), and 4) ability of walking without using aids. All participants provided written informed consent before participation in the study. The university ethics committee approved the experimental procedure.

**Table 1:** Mean (SD) characteristics of the seven participants

| Sex | Age<br>(years) | Height (cm) | Mass (kg) | Preferred walking speed (km/h) | Amputation |  |  |
| --- | --- | --- | --- | --- | --- | --- | --- |
|  |  |  |  |  | Limb | Reason | Time post<br>amputation<br>(years) |
| Male | 45.4<br>(4.2) | 175.7<br>(9.06) | 83.0<br>(5.88) | 2.7 (0.37) | 5<br>right/<br>2 left | Trauma | 24.3 (4.7) |

### 2.2. Marker placement

Fifteen passive reflective markers (14 mm diameter) were attached to the skin of each participant at the right and left bony landmark on the second metatarsal head (toe), calcaneus (heel), lateral malleolus (ankle), mid-tibia, lateral epicondyle of knee (knee), midthigh, anterior superior iliac spine and also on the sacrum, midway between posterior superior iliac spines (S1). Nevertheless, only toe, heel and S1 markers were used in this study.

### 2.3. Task

Before starting the experiment, participants were given enough time to familiarize themselves with walking on the treadmill. During the actual tests, all participants walked on a motorized treadmill (Cosmed^®^ T150, Rome, Italy) at their preferred walking speeds (PWS) in the following four conditions in random order. PWS was recorded following a top-down and bottom-up approach similar to protocols described in previous work by Dingwell and Marin (2006). The average PWS for all participants was 2.70±0.37 km/h. The four walking conditions were: 1) walking with PWS without any video feedback (NVF), 2) walking with PWS with video feedback showing a front view of participant’s walking (FVF), 3) walking with PWS with video feedback showing a back view of participant’s walking (BVF) and, 4) walking with PWS with video feedback showing a lateral view of participant’s walking (LVF). A 42-inch LCD screen was placed in front of the treadmill at the approximate height of the participants to display the full view of the participants’ walking, which was concurrently being recorded by a digital camera and real-time streamed to the screen. Since proper foot placement contribute to avoid instability (McGeer, 1990), participants were instructed to focus on their stepping on the screen while walking normally. To comply with the aim of the study and to avoid providing any extra feedback, no other specific instructions (e.g. how to move their legs, trunk or center of mass) were provided. In addition, participants were not allowed to lean their trunk or head forward while looking at the screen; otherwise the test would be stopped and re-started. Before starting the data recording, participants walked for 1 minute in real test conditions after which data capturing has been started. Participants’ walking was secured with a safety harness without any weight support. Moreover, for safety reasons, an emergency stop band was attached to the participants. Each participant was asked to walk for 2 minutes in each condition. Sufficient rest periods were allocated between the tests to allow participants to recover. All participants walked with their own prostheses and shoes.

### 2.4. Data recording

The three-dimensional coordinate data of the markers were recorded using five Vicon^®^ VCAM motion capture calibrated cameras (Oxford Metrics, Oxford, UK) at the sampling frequency of 100 Hz. Reconstruction and labeling were performed using Vicon^®^ Workstation software (Oxford Metrics, Oxford, UK).

### 2.5. Data analysis

Each step was determined from point of heel contact to the consecutive heel contact of the contralateral foot. Heel contacts were identified as the minima in the heel marker vertical time series (Li, van den Bogert, Caldwell, van Emmerik, & Hamill, 1999). Step length and width were calculated as the distance between heel markers in anterior-posterior (AP) and medio-lateral (ML) directions respectively at heel contact times. Since previous studies (Arshi, Mehdizadeh, & Davids) have shown that linear measure could lead to contradictory results, nonlinear measures were used to quantify foot placement pattern and dynamic stability which will be detailed in the followings.

#### 2.5.1. Quantifying foot placement regularity

Sample entropy (SaEn), a nonlinear measure of time series regularity, was calculated to quantify the effect of video feedback on stepping regularity. Details on the calculation of SaEn already exist in the literature (Lamoth et al., 2010). SaEn measures the degree of regularity or predictability of a time series (Lamoth et al., 2010). It is defined as the probability that a sequence of data points, having repeated itself within a tolerance *r* for a window length *m*, will also repeat itself for *m*+1 points, without allowing self-matches (Lamoth, van Lummel, & Beek, 2009). Smaller SaEn values indicate greater regularity and predictability of the time series. Greater regularity of the kinematic time series in human movement has been reported to be related to higher stability (Lamoth et al., 2010). To calculate SaEn, two input parameters *m*, the window length that will be compared, and *r*, the similarity criterion, were needed. To determine these parameters, the approach suggested in the study of Yentes et al. (2012) was applied. SaEn was calculated for both step length and step width. Since filtering might lead to loss of information at critical points in the time series, non-filtered time-series were used to calculate SaEn (Dingwell & Marin, 2006).

#### 2.5.2. Quantifying symmetry

The symmetry of right and left step lengths was determined using symmetry angle algorithm (Zifchock, Davis, Higginson, & Royer, 2008). Symmetry angle is calculated as the angle of the vector plotted from the right and left step length and width with respect to right horizontal axis. This method reveals a number between 0 and 1, where 0 is associated with complete symmetry. Step length and width were determined as the distance between the right and left heel at heel strike moments in the AP and ML directions, respectively (Nagano, Begg, Sparrow, & Taylor, 2013).

#### 2.5.3. Statistical analyses

Separate one-way repeated measure analyses of variance (ANOVA) were performed to determine the effect of test condition (NVF, FVF, BVF, LVF) on independent measure of SaEn and SA. Post hoc pair-wise comparisons were performed if significant effects were found. Statistical significance was set at *P*<0.05. Data associated with step length and step width were analyzed separately.

## 3. RESULTS

The visual feedback resulted in significant difference of SaEn (Table 2 and Figure 1a) for step length (*P*=0.02) but not step width (*P*=0.73). The pairwise comparisons revealed that the SaEn increased in FVF compared to NVF (*P*=0.04). None of the other conditions affect step length’s SaEn (*P*>0.05). Furthermore, for symmetry angle, the results showed that there was significant effect of visual feedback on symmetry of step length (*P*=0.003; Table 3 and Figure 1b), but not step width (*P*=0.72). Pairwise comparisons indicated a significant difference between NVF and all other conditions (*P*<0.02) except LVF (*P*=0.53) for step length.

**Figure 1:**
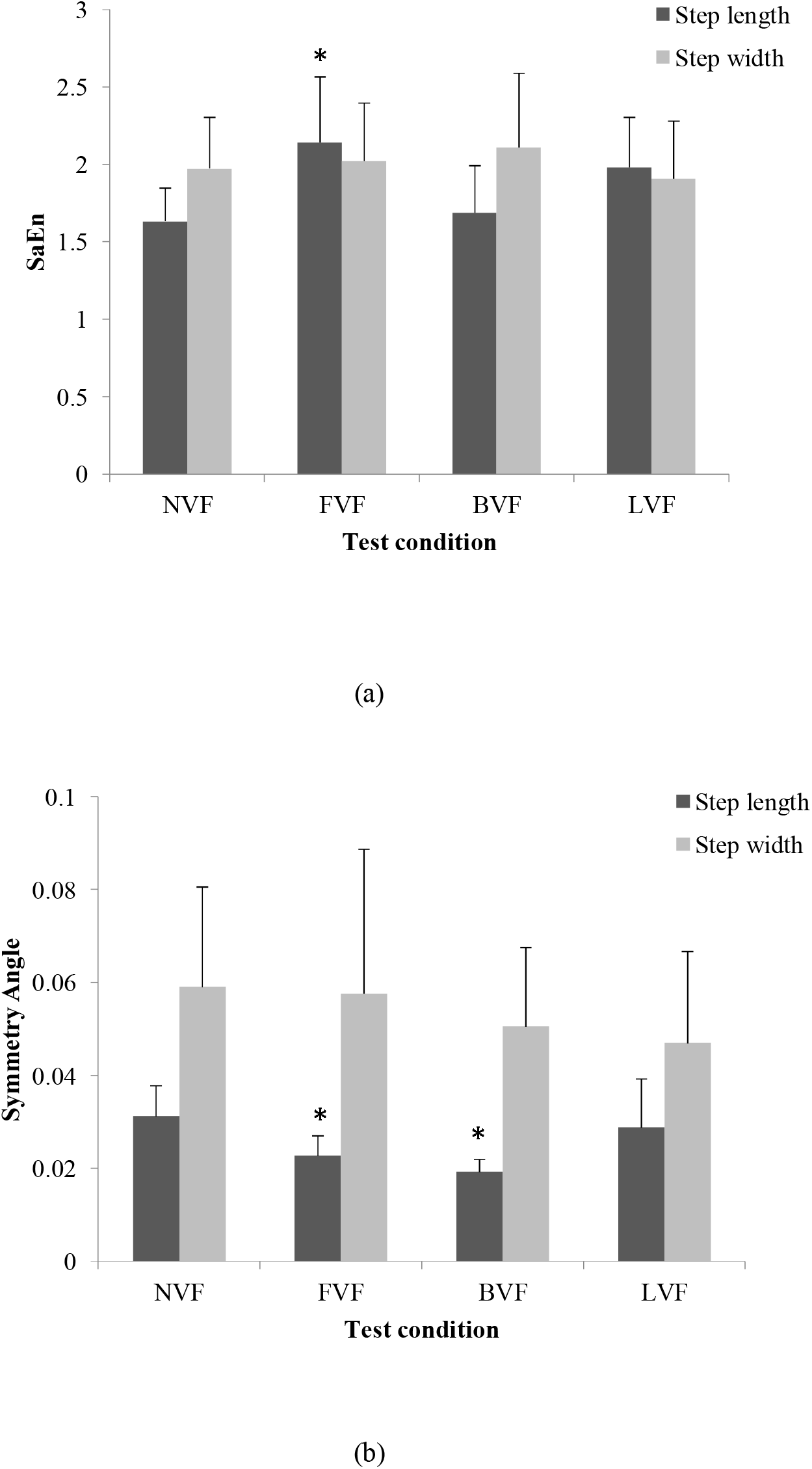
Results of (a) sample entropy (SaEn), and (b) symmetry angle in different test conditions. Error bars indicate standard deviation. Asterisks show significant difference with NVF. NVF= No visual feedback, FVF= Front view visual feedback, BVF= Back view visual feedback, LVF= Lateral view visual feedback.

**Table 2:** results of one-way repeated measure ANOVA test for sample entropy (SaEn) in different test conditions.

|  | Mean (SD) |  |  |  | ANOVA |  |  |
| --- | --- | --- | --- | --- | --- | --- | --- |
| | NVF | FVF | BVF | LVF | F | P-value | $\eta^2$ |
| **Step length** | 1.63 (0.21) | 2.14<br>(0.42) | 1.68<br>(0.30) | 1.98<br>(0.32) | 3.63 | 0.02 | 0.32 |
| **Step width** | 1.97 (0.33) | 2.02<br>(0.37) | 2.10<br>(0.47) | 1.90<br>(0.37) | 0.43 | 0.73 | 0.06 |

$\eta^2$ = effect size (partial eta-squared). NVF= No visual feedback, FVF= Front view visual feedback, BVF= Back view visual feedback, LVF= Lateral view visual feedback

**Table 3:** results of one-way repeated measure ANOVA test for symmetry angle in different test conditions.

|  | Mean (SD) |  |  |  | ANOVA |  |  |
| --- | --- | --- | --- | --- | --- | --- | --- |
| | NVF | FVF | BVF | LVF | F | P-value | $\eta^2$ |
| **Step length** | 0.031<br>(0.006) | 0.022<br>(0.004) | 0.019<br>(0.002) | 0.028<br>(0.010) | 6.98 | 0.003 | 0.53 |
| **Step Width** | 0.059<br>(0.021) | 0.057<br>(0.031) | 0.050<br>(0.017) | 0.046<br>(0.019) | 0.43 | 0.72 | 0.07 |

$\eta^2$ = effect size (partial eta-squared). NVF= No visual feedback, FVF= Front view visual feedback, BVF= Back view visual feedback, LVF= Lateral view visual feedback

## 4. DISCUSSION

The objective of this case study was to determine the feasibility and effectiveness of a real-time video feedback of foot placement on foot placement regularity and symmetry of trans-tibial amputees. The results demonstrated that visual feedback resulted in more regular and also more symmetrical step length in walking of trans-tibial amputees.

The main function of the prescription of a lower limb prosthesis is to increase stability and functionality of the amputees. Since gait temporal-spatial characteristics reflect different locomotor control strategies adopted by individuals during walking, a better understanding of temporal-spatial characteristics of the amputees’ walking could indicate the effect of prosthetic device on their walking. This, as a result, could provide useful information to manage prosthesis prescription and also the rehabilitation process (Parker et al., 2013).

Our results indicated that providing visual feedback resulted in more regular step length (Table 2). In a recent study, Parker et al. (2013) showed that non-faller amputees had more regular AP stepping. Therefore, it could be inferred that increased AP step regularity could contribute to more stable walking. Considering together, adopting more regular step length under the influence of visual feedback by the participants of our study is indicative of more stable gait in AP direction. Nevertheless, our results showed that visual feedback did not affect step width regularity. This indicates that visual feedback from neither of directions could influence gait stability in ML direction. A recent study also reported minimal change in gait stability under perturbations induced in ML direction (Sturdy, Gatesa, Dartera, & Wilkena, 2014). In addition, the participants in our study walked on a motorized treadmill the width of which could restrict their step width adjustment ability. However, this issue needs further investigations with for example participants walking over-ground rather than on a treadmill.

Moreover, our findings indicated that video feedback resulted in significantly more symmetric step length (Table 3 and Figure 1b). This shows that participants were able to adjust their step lengths while watching their walking. Since gait asymmetry may lead to various neuromusculoskeletal problems (Kaufman et al., 2012), the clinical implication of our result is that video feedback gait training could have positive effects on reducing such neuromusculoskeletal problems. More specifically, amputees’ more symmetric step length implies reduction of ground reaction force overload during stance phase of the gait cycle which is found in asymmetrical gaits of amputees (Dingwell, Davis, & Frazier, 1996). The reduction of mechanical overload could thus prevent chronic abnormalities in amputees (Soares, Yamaguti, Mochizuki, Amadio, & Serrão, 2009). Furthermore, previous studies have argued that due to disruption of motor and proprioceptive functions, people with amputation rely more on visual feedback in walking (Beurskens et al., 2014). Our results imply that this reliance on visual feedback was important for amputees to have more symmetrical gait in AP direction. However, this was not the case for step width symmetry. This again might be due to relatively narrow width of the treadmill which makes it difficult to adjust step width while walking on a treadmill.

In a recent study, Hak, van Dieën, van der Wurff, and Houdijk (2014) have suggested that step length asymmetry in people with trans-tibial amputation might be considered as a functional compensation to maintain stability. Our results do not support this argument. That is, in our study, participants’ step length became both more symmetric and more regular (i.e. stable) as a result of the visual feedback. One possible reason for the inconsistency between our results and the results of Hak et al. (2014) study could be associated with adopting different methodologies in quantifying stability. That is, while we used nonlinear measures of SaEn, they used margin of stability to quantify dynamic stability.

Finally, the results of this study showed dissimilar effects of video feedback from different views. Therefore, no general conclusions could be made from our findings on video feedback direction. More investigation adopting eye tracker systems, which determine individuals’ focus of attention, could be helpful in this manner.

One limitation of this study was the low number of participants. Although the number of participants took part in this study (n=7) was in the range of previous studies (e.g. (Beurskens et al., 2014; Lamoth et al., 2010)), a study with greater number of participants would provide stronger statistical results. Furthermore, the findings of this study is associated with treadmill walking which differs to overground walking (Yang & King, 2016). However, the learning effect of this video feedback training might or might not be transferable to over-ground running a concept which requires further investigations.

In conclusion, our findings demonstrated that the video feedback resulted in more symmetrical and more stable step length. These findings demonstrates the feasibility of using a real-time video feedback in gait training of individuals with trans-tibial amputation. This could have positive effects at least in two ways, i) in reducing neuromusculoskeletal problems resulted from asymmetrical and unstable gait, and ii) providing useful information to manage prosthesis prescription. Moreover, these video feedback systems are easy to implement and also affordable which could be adopted by physical therapist in small or medium size clinics.

## Acknowledgments

The tests conducted for this study performed at the Biomechanics Lab, University of Social Welfare and Rehabilitation Sciences.

